# TWEAK and Fn14 expression in human osteoarthritis: evidence for TWEAK-induced RANKL release from chondrocytes

**DOI:** 10.64898/2025.11.30.691475

**Authors:** Anak ASSK Dharmapatni, Tania N Crotti, Renee T Ormsby, Masakazu Kogawa, L. Bogdan Solomon, David M Findlay, David R Haynes, Gerald J Atkins

## Abstract

**Objective:** A role for TWEAK expression and that of its receptor Fn14 by osteoblasts and synovial tissue has been proposed in the pathogenesis of osteoarthritis (OA). Here, we examined whether the cartilage in OA was also a source and target of TWEAK.

**Design:** Articular cartilage samples from 24 patients undergoing hip or knee replacement surgery for OA were investigated for TWEAK and Fn14 expression by both immunostaining and real-time RT-PCR. Samples were graded according to the Pritzker scale from grade 0 (normal) through increasing levels of cartilage damage (grades 1-3). Human primary chondrocytes isolated from OA cartilage were treated with combinations of recombinant TWEAK and TNF and examined for acute effects on *TWEAK*, *FN14*, *RANKL* and *ADAM17* mRNA levels. Released soluble RANKL levels were measured by ELISA.

**Results:** Immunostaining indicated that the majority (21/24) of OA cartilage samples expressed low levels of TWEAK protein and high levels of Fn14, however, expression did not appear to vary with respect to OA grade. *TWEAK* mRNA levels were elevated in grade 2 OA cartilage compared to grade 0 (p < 0.05), while *Fn14* mRNA levels appeared elevated across all damage grades, significantly in grade 2 samples. Primary chondrocytes treated with TNF, alone or in combination with TWEAK, exhibited early upregulated *Fn14* mRNA expression. TWEAK induced *RANKL* mRNA expression. while both TWEAK and TNF induced the mRNA expression of the RANKL sheddase, *ADAM17*. Consistent with these findings, TWEAK and TNF both induced soluble RANKL release by chondrocytes.

**Conclusions:** The expression and regulation of TWEAK and Fn14 is a feature of human OA articular cartilage, suggesting a pathogenetic role. TNF may prime chondrocytes for TWEAK reactivity by upregulating Fn14 expression. The TWEAK and TNF induction of ADAM17 expression and sRANKL release by chondrocytes points to a further disease pathway in human OA.

## Introduction

Osteoarthritis (OA) is the most common human joint pathology, particularly affecting individuals over 65 years of age and is an increasing socioeconomic burden in Western societies due to its prevalence, chronicity and lack of preventive measures or cures, and is the most common reason for joint replacement surgery [1, 2]. OA is characterised by the degeneration of articular cartilage often associated with joint space narrowing, with inadequate compensatory repair of the cartilage, and remodelling and sclerosis of the subchondral bone [3, 4]. The intimate relationship between events in articular cartilage and subchondral bone in OA has been reported elsewhere [3, 5–7].

Receptor activator of NF kappa B ligand (RANKL/TNFSF11), a crucial factor required for osteoclastogenesis, is reported to be expressed by various cell types, including osteoblasts, synovial fibroblasts, T lymphocytes [8–10], osteocytes [11, 12] and chondrocytes [13–15]. RANKL, in both membrane-bound and soluble forms, binds to RANK (TNFRSF11A), which is expressed by osteoclast precursors [16], to induce osteoclastogenesis. Binding of RANKL to the decoy receptor, osteoprotegerin (OPG/TNFRSF11B), inhibits osteoclastogenesis. Several enzymes capable of cleaving cell membrane-bound RANKL and releasing soluble RANKL (sRANKL), termed RANKL sheddases, have been described, including MT1-MMP/MMP14, ADAM10 [17], cathepsin G [18], and TACE/ADAM17 [19, 20]. ADAM17 can also act as a TNF-sheddase [21] and its expression has been shown to be upregulated in human OA cartilage [22]. sRANKL was shown to diffuse into the subchondral bone in an adjuvant induced arthritis (AIA) rabbit model [23]. We and others have demonstrated previously that *RANKL* and *OPG* mRNA levels were altered in human OA cartilage, particularly Pritzker grade 2, with an increased *RANKL*:*OPG* mRNA ratio [24, 25]. The relationship between the *RANKL*:*OPG* mRNA ratio and bone remodelling indices is disrupted in human OA bone [26], and a lower ratio of OPG:RANKL has been reported in OA subchondral bone osteoblasts [27], together supporting a role for RANKL in the aberrant remodelling of OA subchondral bone.

Despite the above evidence, the involvement of the RANKL-OPG axis in human OA aetiology is not fully elucidated, with the current evidence suggesting a role particularly in early OA pathogenesis [28], although other factors including additional TNF family members are clearly also important [29–31]. Previous *in vivo* studies in OA animal models and humans, as well as *in vitro* studies, have demonstrated that chondrocytes produce inflammatory mediators, including TNF and IL-1β, which in turn induce secondary messenger molecules, such as nitric oxide (NO) and prostaglandins and the production of matrix degrading proteases [30, 32].

Another TNF superfamily member, TWEAK (TNFSF12), which signals via the TNF receptor superfamily member, fibroblast growth factor-inducible molecule 14 (Fn14; TNFRSF12), is a mediator with key physiological roles, including tissue repair and wound healing [33–37]. Pathological TWEAK/Fn14 signalling is associated with inflammatory and autoimmune disorders [38, 39]. CD163, expressed by monocytes, acts as a scavenger receptor for soluble TWEAK, potentially ameliorating its proinflammatory effect [40, 41]. In the context of the musculoskeletal system, TWEAK/Fn14 signalling is directly implicated in the joint inflammation and bone erosion associated with rheumatoid arthritis (RA) [42–45]. Circulating TWEAK levels were also elevated in patients with multiple myeloma and correlated with levels of the bone degradation marker, β-CTX [46]. TWEAK has been shown to induce the expression of key osteoblast and osteocyte-associated mediators, including RANKL mRNA and cell surface protein expression by osteoblasts [44, 46], as well as sclerostin [42, 47] and fibroblast growth factor (FGF23) [48] release by osteocytes. We reported previously that TWEAK is expressed in OA synovial tissues and that soluble TWEAK is detectable in OA synovial fluids [44].

In this study, we hypothesised that TWEAK may induce RANKL expression and secretion by OA chondrocytes and hence contribute to the subchondral bone attrition evident in OA. We investigated the expression of TWEAK and Fn14 in various grades of OA cartilage. The effects of recombinant TWEAK and TNF were examined on human chondrocyte expression and release of sRANKL, as well as the expression of related mediators.

## Materials and Methods

### Tissue Samples

Cartilage tissues were obtained with written informed consent from 24 patients undergoing primary knee or hip replacement surgery at the Royal Adelaide Hospital following a diagnosis of advanced osteoarthritis. Cartilage specimens with areas of apparently normal articular cartilage provided the Grade 0 specimens. Clinical data of patients, from whom samples from various grades of OA cartilage tissues were obtained for real time RT-PCR and immunohistochemistry (IHC), are shown in **Table 1**. Cartilage samples were either immediately snap frozen in liquid nitrogen after addition of embedding medium OCT compound (ProScitech, Qld, Australia) for RNA isolation (see below) or immersed in 10% normal buffered formalin overnight before being embedded in paraffin blocks. All procedures and use of human tissues was approved by the Human Research Ethics Committee of the Royal Adelaide Hospital (RAH HREC Approval ID 130114).

**Table 1:** Demographic data of patients used for RT PCR and IHC study.

|  | *Grade 0* | *Grade 1* | *Grade 2* | *Grade 3* |
| --- | --- | --- | --- | --- |
| *Number of samples* | 6 | 6 | 6 | 6 |
| *Mean Age (SEM) (years)* | 88.0 (1.154) | 66.5 (5.92) | 65.2 (6.50) | 71.0 (3.02) |
| *Gender M/F* | 3/3 | 1/5 | 3/3 | 2/4 |
| *Surgery type* | 3 TKR, 3 THR | 4 TKR, 2 THR | 5 TKR, 1 THR | 5 TKR, 1 THR |

TKR: total knee replacement, THR: total hip replacement

### Tissue preparation for Safranin-O staining

Paraffin embedded tissues were cut into 5 µm sections and placed on 3-aminopropyltriethoxy Saline (APTS) coated slides for routine HE and Safranin-O staining for proteoglycans and for immunohistochemical (IHC) detection procedures. Sections were dewaxed in two changes of histolene and two changes of 100% EtOH and were immersed in haematoxylin for 5 min and then 1.5% Safranin-O stain for 6 min. After washing, the slides were then immersed in 0.02% alcoholic fast green solution for 30s and dipped into 1% acetic acid solution. After subsequent washes, the slides were then immersed in 95% alcohol and then 100% alcohol before being mounted with DPX. The cartilage tissue was graded by two blinded observers based on histopathological grading by Pritzker *et al.*[49]. This grading system divides OA tissue into grades 0-6, where grades 0-3 involves only the articular cartilage features, while grades 4-6 also includes the subchondral bone. Since the samples used in this study only contained cartilage, they were graded from 0-3.

### RNA Isolation and generation of cDNA

Thirty-three cartilage samples taken from 24 patients were selected for RT-PCR analysis (grade 0, n = 6, grade 1, n = 9, grade 2, n = 9, grade 3, n = 9). OCT-embedded samples were each cut into five 20 μm sections using a cryostat (Leica, CM1850) and placed in 1.5 ml RNase-free reaction tubes containing TRIzol reagent (600 μl; Thermo Fisher Scientific, Scoresby, VIC, Australia). Total RNA was isolated according to the manufacturer’s instructions. One microgram of total RNA from each sample was reverse transcribed to generate cDNA using Superscript III Reverse Transcriptase (Thermo Fisher Scientific), as described previously [50]. For isolation of total RNA from cultured chondrocytes, 300 μl of TRIzol was added to pooled duplicate wells, each containing approximately 2.5 x 10^5^ cells/well, and processed as above.

### Real Time qRT-PCR

Each PCR tube contained 1 µl cDNA, Platinum SYBR Green PCR Super Mix-UDG, 300 nM of each forward and reverse primers, and DEPC-treated water to a total volume of 15 µl. A template-free negative control was included with each run. PCR was carried out on a Rotor Gene 3000 System (Corbett Life Science, NSW, Australia). All samples were analysed in triplicate and the melt curves obtained after each reaction confirmed the amplification efficacy. The cycle threshold (Ct) values for each gene of interest was then subtracted from the Ct values of the housekeeping gene for data normalisation (*18S*). The comparative Ct method ΔΔCT [51] was subsequently applied to calculate relative gene expression. Primers used were *RANKL*: 5′-GCCTGCGCCGCACCA-3′ (forward) and 5′-CTGCTCTGATGTGCTGTGATCC-3′ (reverse)[52]; *TWEAK*: 5′-ATCGCTGTCCGCCCAGGAGC-3′ (forward) and 5′-CTGTCTGGGGATTCAGTTCCG-3′ (reverse); *FN14*: 5′-TTTCTGGCTTTTTGGTCTGG-3′ (forward) and 5′-CTTGTGGTTGGAGGAGCTTG-3′ (reverse) [53]; *ADAM17*/*TACE*: 5′-CCTTTCTGCGAGAGGGAAC-3′ (forward) and 5′-CACCTTGCAGGAGTTGTCAGT-3′ (reverse). Chondrocyte cultures were also characterised for their basal marker expression: collagen type I/*COL1A1*: 5′-AGGGCTCCAACGAGATCGAGATCCG-3′ (forward) and 5′-TACAGGAAGCAGACAGGGCCAACGTCG-3′ (reverse) [54]; collagen type II (*COL2A1*) 5′-TCCAGGTCTTCAGGGAATGC-3′ (forward) and 5′-AGGACCAGGAGGTCCAACTT-3′ (reverse); collagen type X (*COL10A1*) 5′-GGAGTGTTTTACGCTGAACG-3′ (forward) and 5′-ACCTTGCTCTCCTCTTACTGC-3′ (reverse); aggrecan (*ACAN*) 5′-CTGGAAGTCGTGGTGAAAGGC-3′ (forward) and 5′-AGGAAACTCATCCTTGTCTCCATAGC-3′ (reverse). Housekeeping gene: *18S*: 5′-GCGTTGATTAAGTCCCTGCC-3′ (forward) and 5′-CACCTACGGAAACCTTGTTACGAC-3′ (reverse) [55].

### Immunohistochemical Detection

Formalin fixed paraffin embedded tissues from a subgroup of the patients included above were chosen for immunohistochemical staining of TWEAK and Fn14, essentially as previously described [44]. The primary antibodies included mouse IgG_1_ monoclonal anti-human TWEAK/TNFSF12 (clone P2D10, a gift from Dr Timothy Zheng, Biogen IDEC, Boston MA, USA), and purified anti-human Fn14/TWEAK Receptor/CD266 (314002, clone ITEM-1, BioLegend, San Diego, CA, USA) [43].Antigen retrieval was performed using Proteinase K (50 µg/ml) digestion at 37°C for 30 min. Endogenous peroxidase was blocked using 0.3% v/v hydrogen peroxide in methanol solution for 10 min. After washing with phosphate buffer saline (PBS), blocking serum (provided in Vectastain Universal Elite ABC kit, PK-6200; Vector Laboratories, CA, USA) was applied for 40 min. Primary antibodies anti-TWEAK (15 µg/ml) or anti-Fn14 (10 µg/ml) were then added and incubated in a humidified chamber overnight. Negative controls included omission of primary antibodies and use of non-binding IgG_1_ isotype control antibody (Sigma). Sections were then washed 3 times in PBS and incubated with the secondary antibody, HRP-conjugated biotinylated horse anti-mouse IgG for 45 min. Sections were then treated with Avidin-Biotin Complex reagent for a further 45 min and stained with 3-amino-9-ethylcarbazole (AEC) dye (Cat. No. K3469; DakoCytomation, Glostrup, Denmark) for TWEAK immunostaining or diaminobenzidine (DAB, SK4100, Vector Laboratories, CA, USA) for Fn14 immunostaining. Counterstaining was performed with haematoxylin and lithium carbonate [44].

### In vitro chondrocyte culture

Chondrocytes were isolated, as previously described [56], from fresh cartilage obtained from 5 patients with OA of the knee undergoing total knee replacement surgery. Briefly, articular cartilage was excised from the subchondral bone and placed in serum-free Dulbecco’s Modified Eagle’s Medium (DMEM, Invitrogen Life Technologies, Carlsbad, CA, USA) followed by an overnight digestion in 0.05% w/v bacterial collagenase type II and 0.05% w/v dispase (Invitrogen) in DMEM. The cell suspension was then strained through a 100 μm pore size cell strainer and then centrifuged at 400 xg for 5 min. The cell pellet was resuspended in DMEM contain 10% v/v FBS, 2 mM L-Glutamine, 5 mg/ml penicillin, 50 U/ml streptomycin and Fungizone (Gibco, Life Technologies). Cells were grown in culture monolayers in 75 cm^2^ tissue culture flasks until confluent and then harvested and plated for subsequent experiments.

Upon confluence, cells were harvested using 0.05% w/v trypsin/0.02% w/v EDTA in PBS and then plated in a 12 well plate at a concentration of 2.5 x 10^5^ cells per well for RNA isolation as described above. At 80% confluence, cells were treated with either human recombinant TWEAK (100 ng/ml, a gift from Dr Timothy Zheng, Biogen Idec, Boston, USA), human recombinant TNF (5 ng/ml; R&D Systems, Minneapolis, MN, USA) or a combination of both cytokines for 6 h. Untreated cells were included as the control. Real-time qRT-PCR was also performed to characterise the expression of known chondrocyte markers, *COL1A1*, *COL2A1*, *COL10A1* and aggrecan (*ACAN*) in the primary cultures, and to investigate the relative expression of *RANKL*, *TWEAK*, *Fn14* and *ADAM17* mRNA following various treatments. Supernatants were collected at the end of the culture for assessment of soluble RANKL by ELISA.

### Soluble RANKL ELISA

Resulting levels of sRANKL in chondrocyte supernatants from the above treatments were assessed using a commercial ELISA kit (Cat. No. E90855Hu, Uscn Life Science Inc., Wuhan, China). Absorbance was measured using an ELISA plate reader (Biotek Powerwave, Winooski, VT, USA) at OD_450nm_. A standard curve generated from the assays using KC4 software (Winooski, VT, USA) was used to calculate sRANKL concentrations.

### Statistical Analysis

All data were tested for normality using the Shapiro-Wilk test in GraphPad Prism (GraphPad Software, Boston, MA, USA). The differences in TWEAK or Fn14 protein expression by IHC using the Kruskall Wallis test. All other datasets were normally distributed and were analysed using one-way analysis of variance (ANOVA) with Tukey’s post-hoc tests. A value for p < 0.05 was taken as significant.

## Results

### Histological grading and TWEAK and Fn14 immunostaining in OA cartilage

The histology of each OA grade characterised by Safranin O staining (**Fig. 1**) was similar to that reported previously[24]. TWEAK immunostaining localised mainly to chondrocytes with some staining of the cartilage matrix (**Fig. 1**). Out of the 24 samples, 3 samples showed no positive staining for TWEAK. Staining was perinuclear with cytoplasmic and perilacunar staining also frequently observed. Positive staining for TWEAK was detected in all layers of the cartilage tissue with the superficial layer showing the strongest staining. Fibrochondroblasts in the superficial layers also stained positively for TWEAK. Statistical analysis of semi-quantitative scoring found that there was no significant difference in TWEAK expression between the different OA grades.

**Figure 1:**
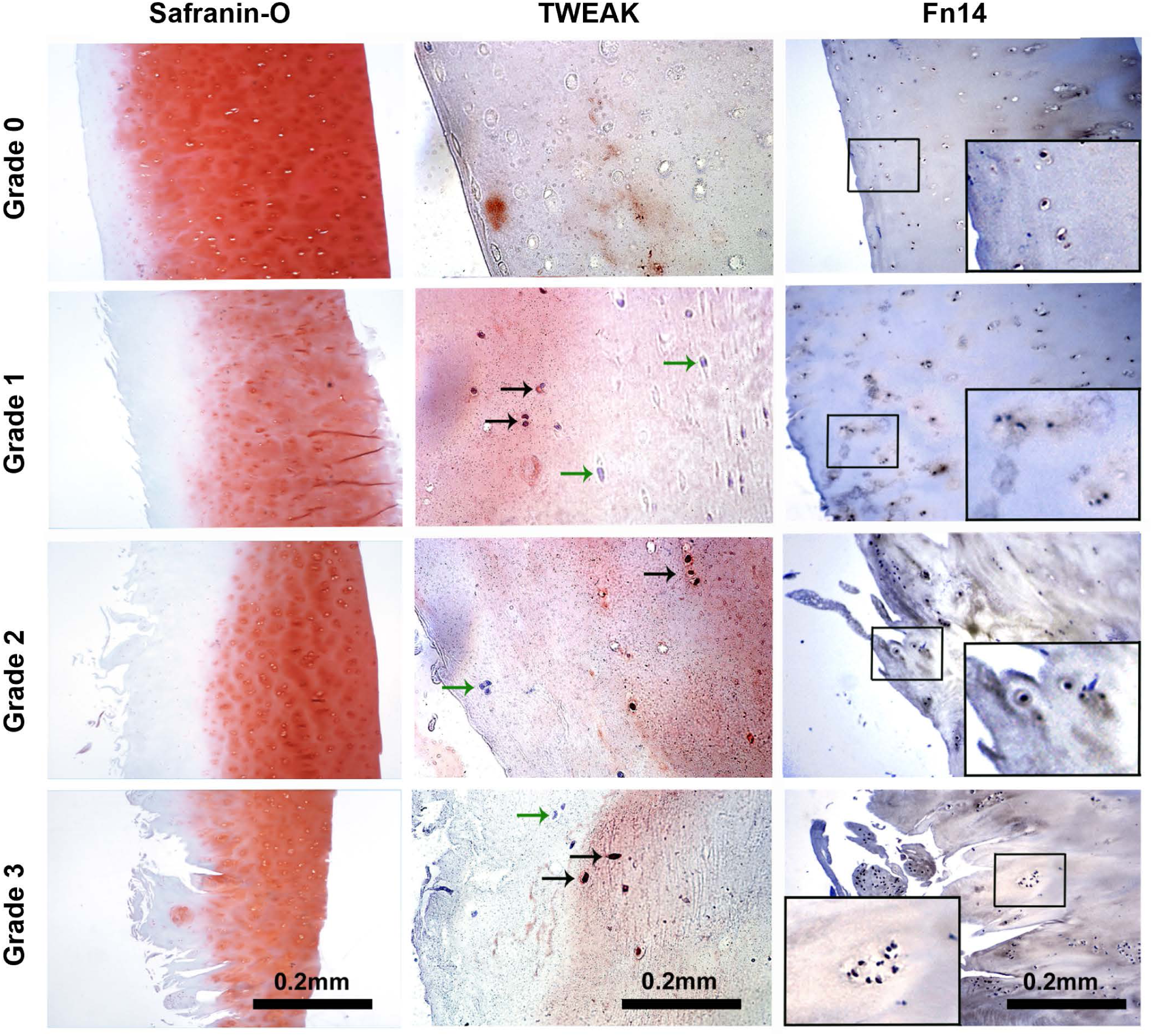
Representative panel showing the histological grading of cartilage samples (grades 0-3) based on Safranin O staining (left column). TWEAK immunostaining (middle column) and FN14 immunostaining (right column) in various grades of OA cartilage. Black arrows indicate examples of TWEAK-positive cells (stained red) and green arrows indicate TWEAK-negative cells. Inserts in Fn14 immunostaining show higher magnification of regions with Fn14-positive cells (stained brown) (magnification x200). Scale bars shown are applicable to all low magnification images.

While TWEAK was expressed by few chondrocytes, Fn14 was expressed by the majority of chondrocytes in all grades of OA cartilage. Positive staining extended into the deep zones of the cartilage with the superficial layers staining more intensely. All cell types present in the samples showed positive staining, including chondrocytes and fibrochondroblasts, with very few negative cells detected (**Fig. 1**). However, semi-quantitative analysis showed that no significant differences existed in overall expression between the OA grades (data not shown).

The relative mRNA expression of *TWEAK* and *Fn14* was assessed in various grades of OA cartilage, comparing expression in grades 1-3 with that seen in grade 0 (**Fig. 2**). Relative *TWEAK* mRNA expression was highest in grade 2 samples and was substantially higher than that of grade 0 (p = 0.0009). Interestingly, expression of *TWEAK* mRNA in grade 2 was also higher compared to grade 3 (p = 0.0035). The relative *Fn14* mRNA expression level was also elevated in grade 2 compared to grade 0 (p = 0.016).

**Figure 2:**
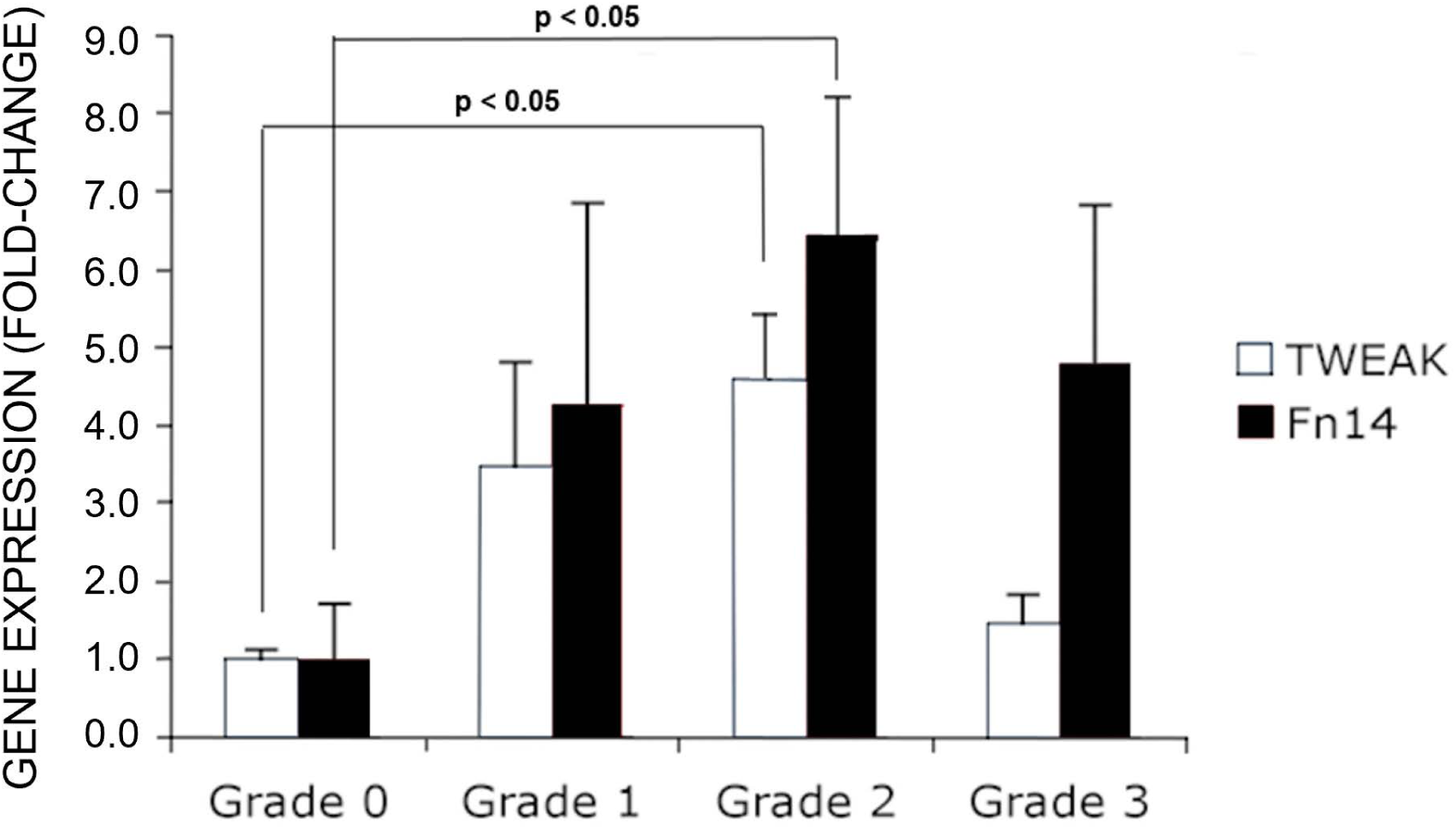
Expression of *TWEAK* and *Fn14* mRNA in various grades of human OA cartilage. Samples were analysed by real-time qRT-PCR, as described in Materials and Methods. Data shown are the mean expression per grade of donors’ tissues (grade 0, *n* = 6, grade 1, *n* = 9, grade 2, *n* = 9, grade 3, *n* = 9) ± standard errors of the mean (SEM). Data were analysed using one-way ANOVA with Tukey’s post-hoc test. Significant differences between grades are indicated by p < 0.05.

### Effect of recombinant TWEAK and TNF on gene expression in chondrocytes

Given the high levels of Fn14 expression in chondrocytes, we next examined the response of human primary chondrocytes to recombinant TWEAK *in vitro*. Cultured chondrocytes isolated from human cartilage expressed collagen types I, II and X, as well as the cartilage matrix gene, aggrecan (**Fig. S1**), confirming their identity as chondrocytes [57]. Chondrocytes from 5 donors were then treated for 6 h with recombinant TWEAK, TNF or a combination of both cytokines (**Fig. 3A-E**). TWEAK mRNA expression was not significantly affected by either treatment (**Fig. 3A**). However, induction of *Fn14* mRNA expression by TNF either alone or in combination with TWEAK was observed (**Fig. 3B**). *RANKL* mRNA induction by TWEAK alone was seen but not with other treatments (**Fig. 3C**). Interestingly, the expression of *ADAM17* mRNA was increased significantly by both TWEAK and TNF (**Fig. 3D**).

**Figure 3:**
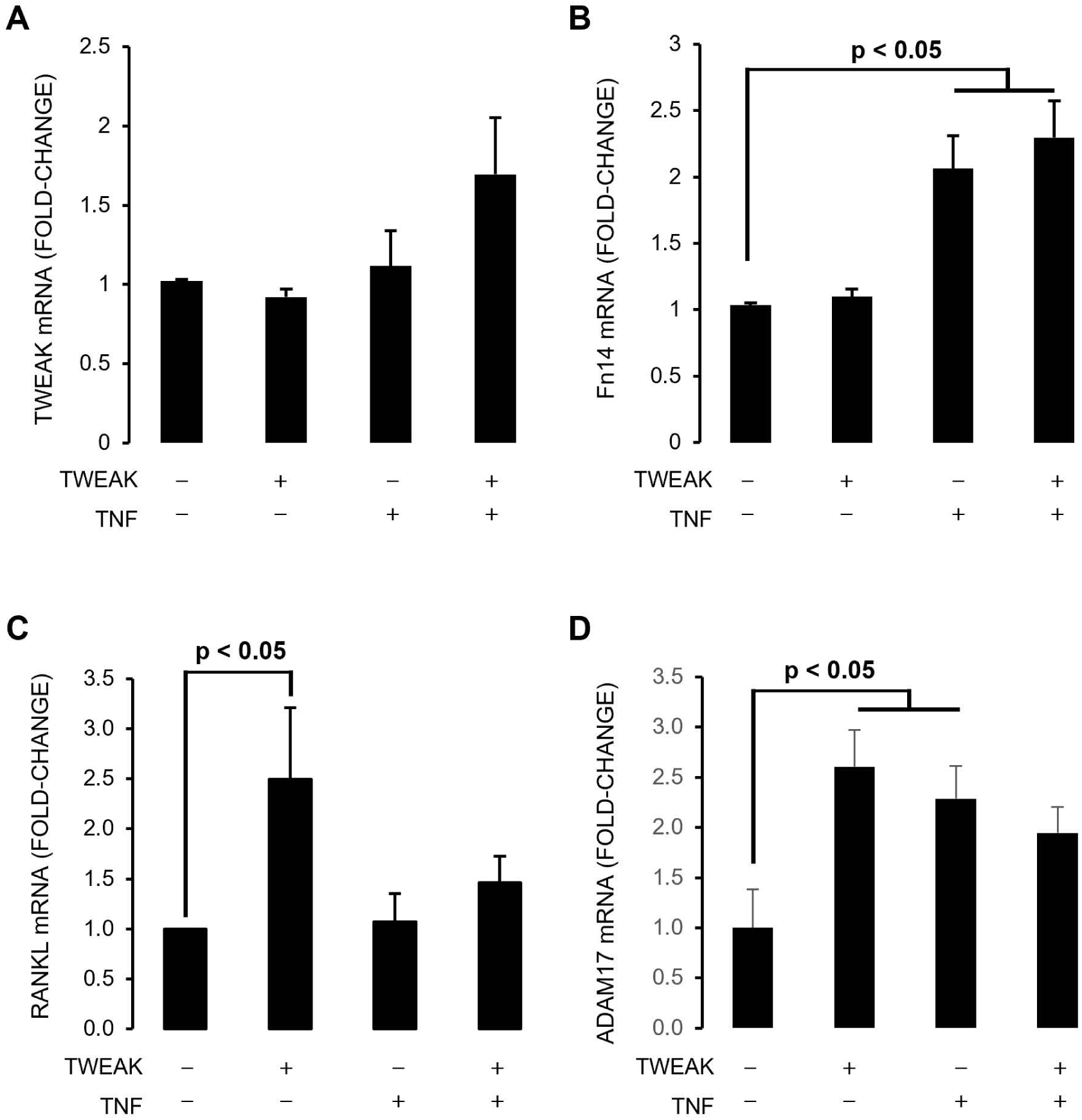
The effect of recombinant TWEAK, recombinant TNF, alone or in combination on gene expression in human primary chondrocyte cultures. Chondrocytes were treated for 6 h and the relative expression of A) *TWEAK*, B) *Fn14*, C) *RANKL* and D) *ADAM17* mRNA determined by real-time qRT-PCR, as described in Materials and Methods. Data shown are means ± SEM pooled from independent experiments using cells derived from individual donors (*n* = 5). Data were analysed by one-way ANOVA with Tukey’s post-hoc test. Significant differences to untreated are indicated by p < 0.05.

### Effect of TWEAK and TNFα on soluble RANKL secretion

Treatment of chondrocytes with TWEAK or TNF significantly increased the levels of sRANKL in culture supernatants after 24 h compared to untreated chondrocytes (p < 0.05), with a combination treatment giving the maximal effect of an approximately 4-fold increase in comparison to untreated chondrocytes (**Fig. 4**).

**Figure 4:**
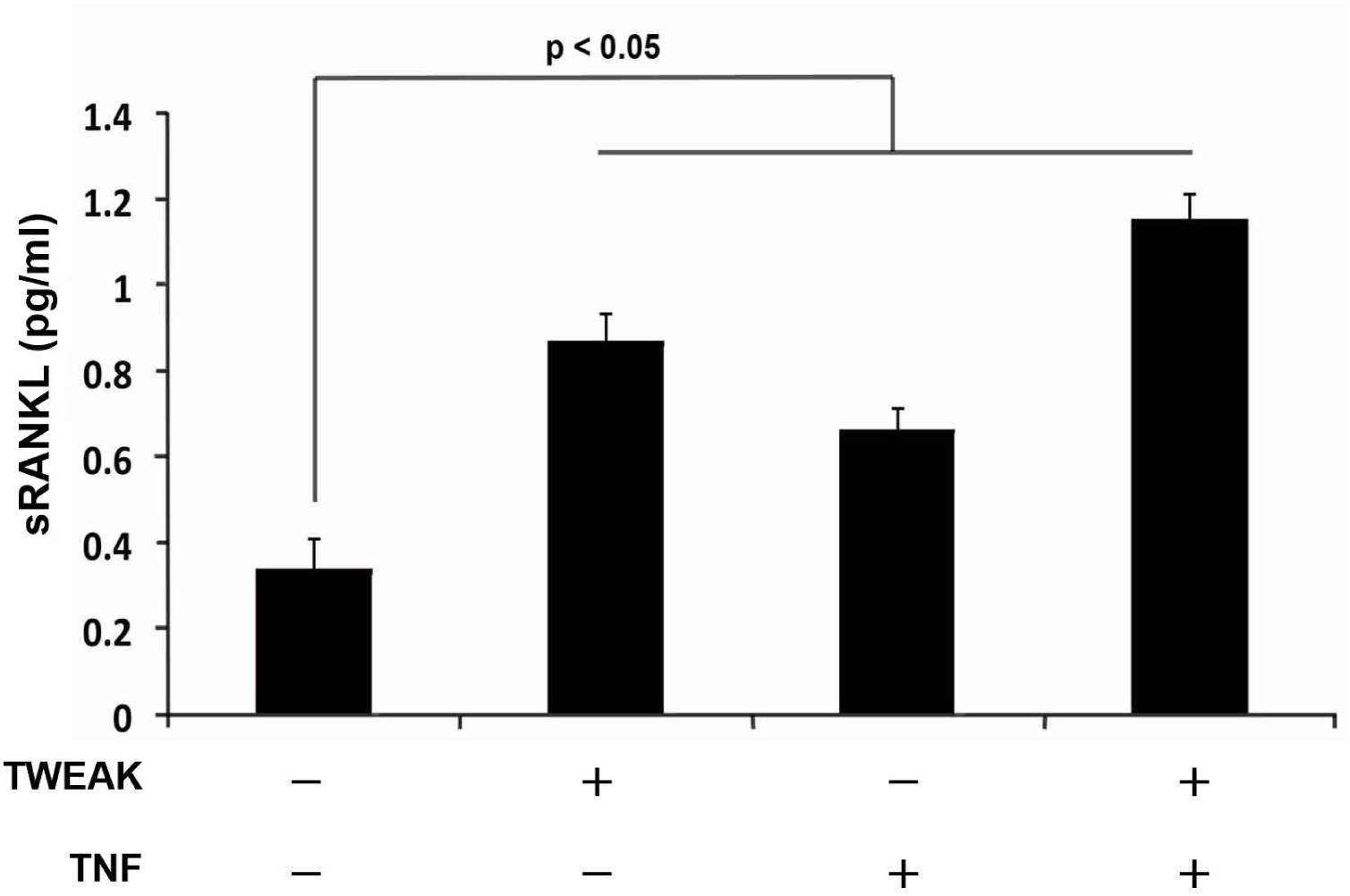
sRANKL levels in chondrocyte culture supernatant following various treatments after 24 h, as assessed by ELISA. Data were analysed using one-way ANOVA with Tukey’s post-hoc test. Data shown are mean levels ± SEM (*n* = 5 donors). Significant differences to untreated culture levels are indicated by p < 0.05.

## Discussion

In this study, we investigated the expression and potential role of TWEAK and its receptor, Fn14, in human OA cartilage. Significantly higher expression of both *TWEAK* and *Fn14* mRNA in grade 2 compared to grade 0 OA cartilage was observed. Differences between various OA grades observed at mRNA levels, however, were not observed at the protein level in terms of the numbers of cells stained. Both membrane bound and soluble TWEAK have been reported in RA and OA [58], which together with results from this and our previous study [44], suggests that both forms are present within the OA joint. It may be that TWEAK protein is efficiently secreted by chondrocytes, perhaps explaining the discrepancy between TWEAK gene expression and immunostaining. Consistent with this possibility, the observed immunostaining in chondrocytes was mostly either perinuclear or associated with the extracellular matrix. Although there was a better correlation between mRNA and protein levels in the case of Fn14, there was no statistically significant difference between the various OA grades, with the majority of chondrocytes staining positive for Fn14 in all grades above grade 0. This was surprising since it has been proposed that Fn14 levels are influenced by the degree of tissue damage rather than inflammation [59]. It is possible that protein levels on a per cell basis varied between OA grades, but this difference could not be discerned by immunostaining.

We have previously reported significantly higher expression of TWEAK and Fn14 in OA, as well as RA synovial tissues compared to synovial tissues from healthy controls [44], with several cell types expressing TWEAK and Fn14 within the OA joint. We have also demonstrated detectable levels of soluble TWEAK in OA synovial fluid (average of 713 ± 134 pg/ml) [44]. These levels are much higher than those reported in the serum of OA patients (47.3 ± 31.0 pg/ml) [60], suggesting that local sources of TWEAK within the OA joint play an important role. Our findings here suggest that chondrocytes contribute to this pool of soluble TWEAK. Detectable levels of TNF have also been previously reported in OA synovial fluid [61, 62]. Consistent with previous reports that TNF can act synergistically with TWEAK [47], TNF upregulated both *TWEAK* and *Fn14* mRNA expression in human chondrocytes *in vitro*, which suggests that the combination of TWEAK and TNF contribute together to the cartilage and bone pathology in OA.

We have also previously published the high expression of *RANKL* mRNA in grade 2 OA cartilage compared to grade 0 [44], in a pattern similar to that observed here for *TWEAK* and *Fn14* mRNA, suggesting that there may be crosstalk between the respective signalling pathways in OA. Similar to our previous findings in osteoblasts [44, 46], we showed that TWEAK induced *RANKL* mRNA expression in primary chondrocytes *in vitro*, while TNF did not. Despite minor effects at the mRNA level, both TWEAK and TNF stimulated soluble RANKL protein release from chondrocytes. As a potential explanation for this, both TWEAK and TNF induced the expression of ADAM17/TACE. ADAM17 has been reported to be upregulated in human OA cartilage [22] and TWEAK was shown to induce ADAM17 expression in renal tubular epithelium [63]. ADAM17 was shown previously to regulate sRANKL release from human primary osteoblasts [19]. Detectable levels of sRANKL in synovial fluid [64] and plasma [65] of patients with OA have also been reported previously. In a rabbit model of OA, it has been reported that sRANKL can diffuse from cartilage to the bone cartilage interface and stimulate osteoclastogenesis, resulting in bone degradation [23]. Taken together, these studies and our current results indicate that sRANKL may be released at the cartilage-bone interface of OA joint in the presence of TWEAK and TNF by a mechanism involving ADAM17-mediated cleavage.

This study has a number of limitations. Firstly, the sample size was relatively small and likely prevented us from discerning significant differences in TWEAK and Fn14 expression within Pritzker grades 1-3, for example. Furthermore, we used disaggregated chondrocytes for *in vitro* gene expression assays under standard cell culture conditions, including a cell expansion step due to the inherently low cell yields from human cartilage. Thus, we cannot rule out that the cells underwent a degree of phenotypic change prior to treatment and the responses in intact cartilage may differ due to native cell-cell matrix interactions and 3-dimensional morphology, as well as the low oxygen (hypoxic) conditions in cartilage *in vivo*. These limitations were mitigated to a certain extent by confirming expression of key chondrogenic markers in the cultured cells used and treating chondrocytes for a short time period (6 hours) to identify proof-of-principle, direct effects of agonists. This approach was additionally used due to the anticipated difficulty in efficiently delivering recombinant cytokine treatments to cells buried deep within a cartilaginous matrix. However, future experiments would likely benefit from the use of a microfluidic-based organ culture system. Finally, OA is recognised as a disease of the whole joint and the analysis of chondrocytes only missed potentially important interactions with other cell types, such as sub-chondral bone cells [28, 66, 67]. Most relevant to the mediators examined here, we have reported the ability of TWEAK and TNF to potently induce the expression of a negative regulator of bone formation, *SOST*/sclerostin by human osteoblasts [47], which may inhibit the repair and/or the mineralisation of subchondral bone in OA [68] and contribute to RANKL expression by osteocytes, increasing osteoclastic activity [69].

## Conclusions

In conclusion, this study indicates that chondrocytes in the articular cartilage in OA are both sources of and targets for TWEAK. Together with our previous study demonstrating that TWEAK is expressed by cells in OA synovial tissue, our findings are supportive of a role for TNF and TWEAK in the pathophysiology of human OA, leading to increased release of sRANKL by chondrocytes, possibly mediated through induction of ADAM17 expression.

## Ethics Statement

All procedures followed were in accordance with the ethical standards of the responsible committees on human experimentation, Central Adelaide Local Health Network (CALHN) and The University of Adelaide Human Research Ethics Committee, and with the Helsinki Declaration of 1975, as revised in 2000. The study received specific human research ethics approval (RAH HREC Approval ID 130114).

## Acknowledgements

The authors are grateful to the surgeons and nursing staff of the Department of Orthopaedics & Trauma, Royal Adelaide Hospital for the provision of human cartilage samples. The authors thank Professor Malcolm Smith for intellectual input and Dr Timothy S Zheng (Biogen Idec, Boston, MA, USA) for provision of antibodies to TWEAK and Fn14 as well as recombinant human TWEAK, and Mr Dale Caville for assisting in photographic and figure presentation.

---

**Supplementary Figure 1:**
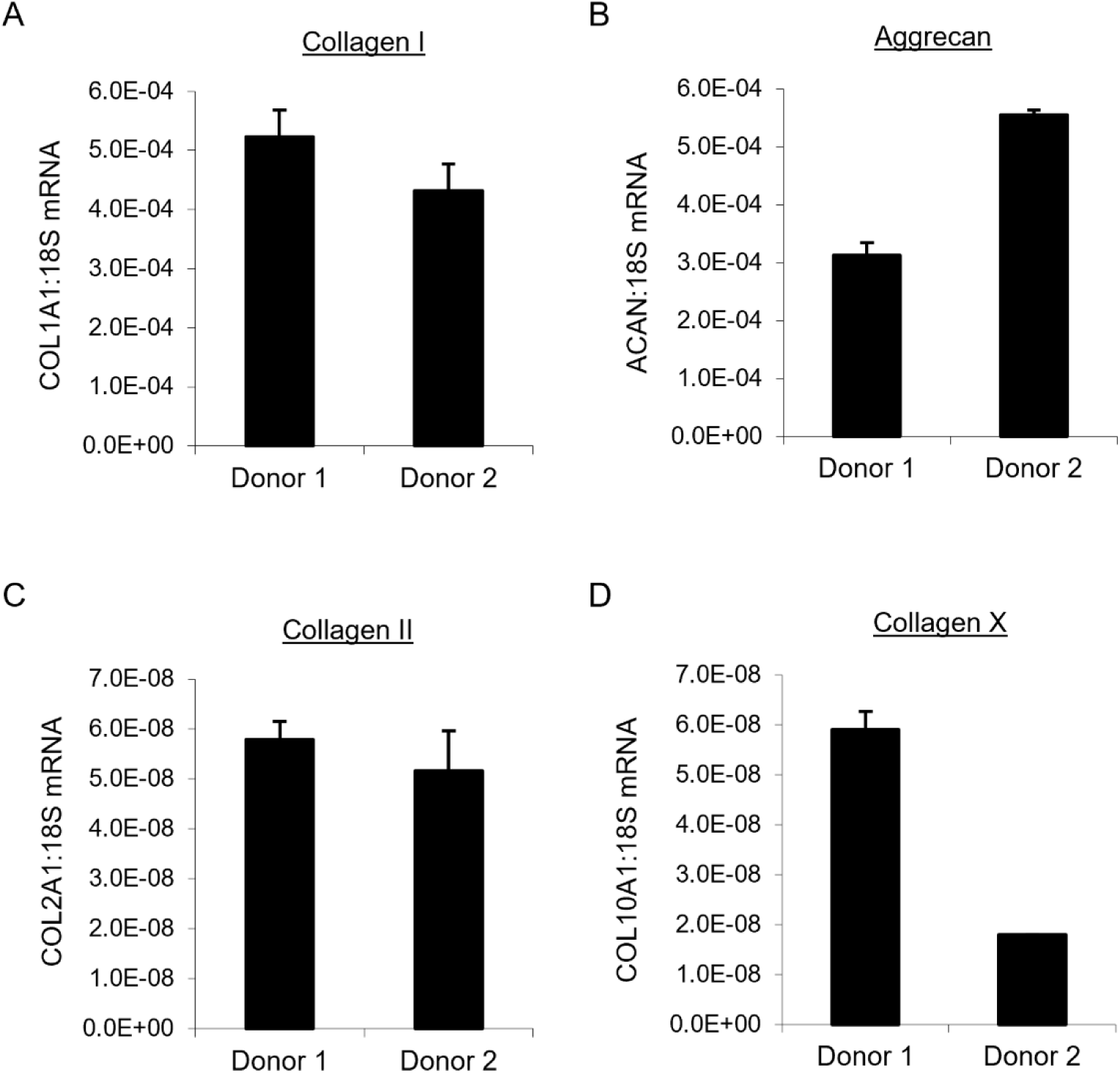
Expression of chondrocyte markers in human chondrocyte cultures *in vitro*, as determined by real-time RT-PCR: A) Type I collagen, B) Aggrecan, C) Type II collagen, D) Type X collagen. Data for 2 donors’ cells are shown as mean expression ± standard deviation of triplicate reactions expressed relative to that of 18S rRNA expression.

## REFERENCES

[1] J.A. Buckwalter, J.A. Martin, Osteoarthritis, Advanced drug delivery reviews 58(2) (2006) 150–67.

[2] D.J. Hunter, S. Bierma-Zeinstra, Osteoarthritis, The Lancet 393(10182) (2019) 1745–1759.

[3] D.D. Kumarasinghe, B. Hopwood, J.S. Kuliwaba, G.J. Atkins, N.L. Fazzalari, An update on primary hip osteoarthritis including altered Wnt and TGF-beta associated gene expression from the bony component of the disease, Rheumatology (Oxford) 50(12) (2011) 2166–75.

[4] J.A. Buckwalter, H.J. Mankin, Articular cartilage: degeneration and osteoarthritis, repair, regeneration, and transplantation, Instructional course lectures 47 (1998) 487–504.

[5] S.K. Tat, J.P. Pelletier, C.R. Velasco, M. Padrines, J. Martel-Pelletier, New perspective in osteoarthritis: the OPG and RANKL system as a potential therapeutic target?, The Keio journal of medicine 58(1) (2009) 29–40.

[6] M.B. Goldring, S.R. Goldring, Articular cartilage and subchondral bone in the pathogenesis of osteoarthritis, Annals of the New York Academy of Sciences 1192 (2010) 230–7.

[7] D.M. Findlay, G.J. Atkins, Osteoblast-Chondrocyte Interactions in Osteoarthritis, Current osteoporosis reports (2014).

[8] H. Yasuda, N. Shima, N. Nakagawa, K. Yamaguchi, M. Kinosaki, S. Mochizuki, A. Tomoyasu, K. Yano, M. Goto, A. Murakami, E. Tsuda, T. Morinaga, K. Higashio, N. Udagawa, N. Takahashi, T. Suda, Osteoclast differentiation factor is a ligand for osteoprotegerin/osteoclastogenesis-inhibitory factor and is identical to TRANCE/RANKL, Proc Natl Acad Sci U S A 95(7) (1998) 3597–602.

[9] N. Udagawa, S. Kotake, N. Kamatani, N. Takahashi, T. Suda, The molecular mechanism of osteoclastogenesis in rheumatoid arthritis, Arthritis research 4(5) (2002) 281–9.

[10] E.M. Gravallese, Bone destruction in arthritis, Annals of the rheumatic diseases 61 Suppl 2 (2002) ii84–6.

[11] J. Xiong, C.A. O’Brien, Osteocyte RANKL: new insights into the control of bone remodeling, J Bone Miner Res 27(3) (2012) 499–505.

[12] M. Prideaux, D.M. Findlay, G.J. Atkins, Osteocytes: The master cells in bone remodelling, Curr Opin Pharmacol 28 (2016) 24–30.

[13] S. Kwan Tat, N. Amiable, J.P. Pelletier, C. Boileau, D. Lajeunesse, N. Duval, J. Martel-Pelletier, Modulation of OPG, RANK and RANKL by human chondrocytes and their implication during osteoarthritis, Rheumatology (Oxford) 48(12) (2009) 1482–90.

[14] V. Marzaioli, J.P. McMorrow, H. Angerer, A. Gilmore, D. Crean, D. Zocco, P. Rooney, D. Veale, U. Fearon, M. Gogarty, A.N. McEvoy, M.H. Stradner, E.P. Murphy, Histamine contributes to increased RANKL to osteoprotegerin ratio through altered nuclear receptor 4A activity in human chondrocytes, Arthritis and rheumatism 64(10) (2012) 3290–301.

[15] A. Takeshita, K. Nishida, A. Yoshida, Y. Nasu, R. Nakahara, D. Kaneda, H. Ohashi, T. Ozaki, RANKL expression in chondrocytes and its promotion by lymphotoxin-α in the course of cartilage destruction during rheumatoid arthritis, PLoS One 16(7) (2021) e0254268.

[16] G.J. Atkins, P. Kostakis, C. Vincent, A.N. Farrugia, J.P. Houchins, D.M. Findlay, A. Evdokiou, A.C. Zannettino, RANK Expression as a cell surface marker of human osteoclast precursors in peripheral blood, bone marrow, and giant cell tumors of bone, J Bone Miner Res 21(9) (2006) 1339–49.

[17] A. Hikita, I. Yana, H. Wakeyama, M. Nakamura, Y. Kadono, Y. Oshima, K. Nakamura, M. Seiki, S. Tanaka, Negative regulation of osteoclastogenesis by ectodomain shedding of receptor activator of NF-kappaB ligand, J Biol Chem 281(48) (2006) 36846–55.

[18] T.J. Wilson, K.C. Nannuru, M. Futakuchi, A. Sadanandam, R.K. Singh, Cathepsin G enhances mammary tumor-induced osteolysis by generating soluble receptor activator of nuclear factor-kappaB ligand, Cancer Res 68(14) (2008) 5803–11.

[19] B. Pan, A.N. Farrugia, L.B. To, D.M. Findlay, J. Green, K. Lynch, A.C. Zannettino, The nitrogen-containing bisphosphonate, zoledronic acid, influences RANKL expression in human osteoblast-like cells by activating TNF-alpha converting enzyme (TACE), J Bone Miner Res 19(1) (2004) 147–54.

[20] L. Lum, B.R. Wong, R. Josien, J.D. Becherer, H. Erdjument-Bromage, J. Schlondorff, P. Tempst, Y. Choi, C.P. Blobel, Evidence for a role of a tumor necrosis factor-alpha (TNF-alpha)-converting enzyme-like protease in shedding of TRANCE, a TNF family member involved in osteoclastogenesis and dendritic cell survival, J Biol Chem 274(19) (1999) 13613–8.

[21] R.A. Black, C.T. Rauch, C.J. Kozlosky, J.J. Peschon, J.L. Slack, M.F. Wolfson, B.J. Castner, K.L. Stocking, P. Reddy, S. Srinivasan, N. Nelson, N. Boiani, K.A. Schooley, M. Gerhart, R. Davis, J.N. Fitzner, R.S. Johnson, R.J. Paxton, C.J. March, D.P. Cerretti, A metalloproteinase disintegrin that releases tumour-necrosis factor-alpha from cells, Nature 385(6618) (1997) 729–33.

[22] I.R. Patel, M.G. Attur, R.N. Patel, S.A. Stuchin, R.A. Abagyan, S.B. Abramson, A.R. Amin, TNF-alpha convertase enzyme from human arthritis-affected cartilage: isolation of cDNA by differential display, expression of the active enzyme, and regulation of TNF-alpha, Journal of immunology (Baltimore, Md. : 1950) 160(9) (1998) 4570–9.

[23] M.J. Martinez-Calatrava, I. Prieto-Potin, J.A. Roman-Blas, L. Tardio, R. Largo, G. Herrero-Beaumont, RANKL synthesized by articular chondrocytes contributes to juxta-articular bone loss in chronic arthritis, Arthritis research & therapy 14(3) (2012) R149.

[24] A.R. Upton, C.A. Holding, A.A. Dharmapatni, D.R. Haynes, The expression of RANKL and OPG in the various grades of osteoarthritic cartilage, Rheumatol Int 32(2) (2012) 535–40.

[25] H. Komuro, T. Olee, K. Kuhn, J. Quach, D.C. Brinson, A. Shikhman, J. Valbracht, L. Creighton-Achermann, M. Lotz, The osteoprotegerin/receptor activator of nuclear factor kappaB/receptor activator of nuclear factor kappaB ligand system in cartilage, Arthritis and rheumatism 44(12) (2001) 2768–76.

[26] N.L. Fazzalari, J.S. Kuliwaba, G.J. Atkins, M.R. Forwood, D.M. Findlay, The ratio of messenger RNA levels of receptor activator of nuclear factor kappaB ligand to osteoprotegerin correlates with bone remodeling indices in normal human cancellous bone but not in osteoarthritis, J Bone Miner Res 16(6) (2001) 1015–27.

[27] S. Kwan Tat, J.P. Pelletier, D. Lajeunesse, H. Fahmi, M. Lavigne, J. Martel-Pelletier, The differential expression of osteoprotegerin (OPG) and receptor activator of nuclear factor kappaB ligand (RANKL) in human osteoarthritic subchondral bone osteoblasts is an indicator of the metabolic state of these disease cells, Clinical and experimental rheumatology 26(2) (2008) 295–304.

[28] D. Muratovic, G.J. Atkins, D.M. Findlay, Is RANKL a potential molecular target in osteoarthritis?, Osteoarthritis Cartilage 32(5) (2024) 493–500.

[29] T. Lawyer, S. Wingerter, M. Tucci, H. Benghuzzi, Cellular effects of catabolic inflammatory cytokines on chondrocytes-biomed 2011, Biomedical sciences instrumentation 47 (2011) 252–7.

[30] J.C. Fernandes, J. Martel-Pelletier, J.P. Pelletier, The role of cytokines in osteoarthritis pathophysiology, Biorheology 39(1-2) (2002) 237–46.

[31] M. Kobayashi, G.R. Squires, A. Mousa, M. Tanzer, D.J. Zukor, J. Antoniou, U. Feige, A.R. Poole, Role of interleukin-1 and tumor necrosis factor alpha in matrix degradation of human osteoarthritic cartilage, Arthritis and rheumatism 52(1) (2005) 128–35.

[32] M.B. Goldring, K.B. Marcu, Cartilage homeostasis in health and rheumatic diseases, Arthritis research & therapy 11(3) (2009) 224.

[33] A.T. Khalaf, X. Liu, Targeting TWEAK/Fn14 pathway: Implications for stem cell regulation and advancements in skin disease therapies, Int Immunopharmacol 167 (2025) 115699.

[34] B. Liu, W. Wu, W. Hu, H. Zheng, T. Lan, Research progress on TWEAK/Fn14 signaling in chronic wound healing, Front Surg 13 (2026) 1769996.

[35] L.C. Burkly, TWEAK/Fn14 axis: the current paradigm of tissue injury-inducible function in the midst of complexities, Semin Immunol 26(3) (2014) 229–36.

[36] J.A. Winkles, The TWEAK-Fn14 cytokine-receptor axis: discovery, biology and therapeutic targeting, Nat Rev Drug Discov 7(5) (2008) 411–25.

[37] M. Bhattacharjee, R. Raju, A. Radhakrishnan, V. Nanjappa, B. Muthusamy, K. Singh, D. Kuppusamy, B.T. Lingala, A. Pan, P.P. Mathur, H.C. Harsha, T.S. Prasad, G.J. Atkins, A. Pandey, A. Chatterjee, A Bioinformatics Resource for TWEAK-Fn14 Signaling Pathway, J Signal Transduct 2012 (2012) 376470.

[38] S. Campbell, J. Michaelson, L. Burkly, C. Putterman, The role of TWEAK/Fn14 in the pathogenesis of inflammation and systemic autoimmunity, Front Biosci 9 (2004) 2273–84.

[39] W.D. Xu, Y. Zhao, Y. Liu, Role of the TWEAK/Fn14 pathway in autoimmune diseases, Immunol Res 64(1) (2016) 44–50.

[40] L.C. Bover, M. Cardo-Vila, A. Kuniyasu, J. Sun, R. Rangel, M. Takeya, B.B. Aggarwal, W. Arap, R. Pasqualini, A previously unrecognized protein-protein interaction between TWEAK and CD163: potential biological implications, Journal of immunology (Baltimore, Md. : 1950) 178(12) (2007) 8183–94.

[41] J.K. Qian, Y. Ma, X. Huang, X.R. Li, Y.F. Xu, Z.Y. Liu, Y. Gu, K. Shen, L.J. Tian, Y.T. Wang, N.N. Cheng, B.S. Yang, K.Y. Huang, Y. Chai, G.Q. Liu, N.Q. Cui, S.Y. Deng, N. Jiang, D.R. Xu, B. Yu, The CD163/TWEAK/Fn14 axis: A potential therapeutic target for alleviating inflammatory bone loss, J Orthop Translat 49 (2024) 82–95.

[42] D.M. Findlay, G.J. Atkins, TWEAK and TNF regulation of sclerostin: a novel pathway for the regulation of bone remodelling, Adv Exp Med Biol 691 (2011) 337–48.

[43] S.J. Perper, B. Browning, L.C. Burkly, S. Weng, C. Gao, K. Giza, L. Su, L. Tarilonte, T. Crowell, L. Rajman, L. Runkel, M. Scott, G.J. Atkins, D.M. Findlay, T.S. Zheng, H. Hess, TWEAK is a novel arthritogenic mediator, Journal of immunology (Baltimore, Md. : 1950) 177(4) (2006) 2610–20.

[44] A.A. Dharmapatni, M.D. Smith, T.N. Crotti, C.A. Holding, C. Vincent, H.M. Weedon, A.C. Zannettino, T.S. Zheng, D.M. Findlay, G.J. Atkins, D.R. Haynes, TWEAK and Fn14 expression in the pathogenesis of joint inflammation and bone erosion in rheumatoid arthritis, Arthritis research & therapy 13(2) (2011) R51.

[45] Y.Y. Du, Y.X. Zhao, Y.P. Liu, W. Liu, M.M. Wang, C.M. Yuan, Regulatory Tweak/Fn14 signaling pathway as a potent target for controlling bone loss, Biomed Pharmacother 70 (2015) 170–3.

[46] S.A. Williams, S.K. Martin, C. Vincent, S. Gronthos, T. Zheng, G.J. Atkins, A.C. Zannettino, Circulating levels of TWEAK correlate with bone erosion in multiple myeloma patients, Br J Haematol 150(3) (2010) 373–6.

[47] C. Vincent, D.M. Findlay, K.J. Welldon, A.R. Wijenayaka, T.S. Zheng, D.R. Haynes, N.L. Fazzalari, A. Evdokiou, G.J. Atkins, Pro-inflammatory cytokines TNF-related weak inducer of apoptosis (TWEAK) and TNFalpha induce the mitogen-activated protein kinase (MAPK)-dependent expression of sclerostin in human osteoblasts, J Bone Miner Res 24(8) (2009) 1434–49.

[48] N. Ito, A.R. Wijenayaka, M. Prideaux, M. Kogawa, R.T. Ormsby, A. Evdokiou, L.F. Bonewald, D.M. Findlay, G.J. Atkins, Regulation of FGF23 expression in IDG-SW3 osteocytes and human bone by pro-inflammatory stimuli, Mol Cell Endocrinol 399 (2015) 208–18.

[49] K.P. Pritzker, S. Gay, S.A. Jimenez, K. Ostergaard, J.P. Pelletier, P.A. Revell, D. Salter, W.B. van den Berg, Osteoarthritis cartilage histopathology: grading and staging, Osteoarthritis Cartilage 14(1) (2006) 13–29.

[50] K.J. Welldon, D.M. Findlay, A. Evdokiou, R.T. Ormsby, G.J. Atkins, Calcium induces pro-anabolic effects on human primary osteoblasts associated with acquisition of mature osteocyte markers, Mol Cell Endocrinol 376(1-2) (2013) 85–92.

[51] K.J. Livak, T.D. Schmittgen, Analysis of relative gene expression data using real-time quantitative PCR and the 2(-Delta Delta C(T)) Method, Methods. 25(4) (2001) 402–8.

[52] C.A. Holding, D.M. Findlay, R. Stamenkov, S.D. Neale, H. Lucas, A.S. Dharmapatni, S.A. Callary, K.R. Shrestha, G.J. Atkins, D.W. Howie, D.R. Haynes, The correlation of RANK, RANKL and TNFalpha expression with bone loss volume and polyethylene wear debris around hip implants, Biomaterials 27(30) (2006) 5212–9.

[53] N.G. Kataria, P.M. Bartold, A.A. Dharmapatni, G.J. Atkins, C.A. Holding, D.R. Haynes, Expression of tumor necrosis factor-like weak inducer of apoptosis (TWEAK) and its receptor, fibroblast growth factor-inducible 14 protein (Fn14), in healthy tissues and in tissues affected by periodontitis, J Periodontal Res 45(4) (2010) 564–73.

[54] G.J. Atkins, K.J. Welldon, C.A. Holding, D.R. Haynes, D.W. Howie, D.M. Findlay, The induction of a catabolic phenotype in human primary osteoblasts and osteocytes by polyethylene particles, Biomaterials 30(22) (2009) 3672–81.

[55] D. Yang, A.R. Wijenayaka, L.B. Solomon, S.M. Pederson, D.M. Findlay, S.P. Kidd, G.J. Atkins, Novel Insights into Staphylococcus aureus Deep Bone Infections: the Involvement of Osteocytes, mBio 9(2) (2018).

[56] D.J. Huey, K.A. Athanasiou, Alteration of the fibrocartilaginous nature of scaffoldless constructs formed from leporine meniscus cells and chondrocytes through manipulation of culture and processing conditions, Cells, tissues, organs 197(5) (2013) 360–71.

[57] A.M. Fernandes, S.R. Herlofsen, T.A. Karlsen, A.M. Kuchler, Y. Floisand, J.E. Brinchmann, Similar properties of chondrocytes from osteoarthritis joints and mesenchymal stem cells from healthy donors for tissue engineering of articular cartilage, PLoS One 8(5) (2013) e62994.

[58] J.S. Park, M.K. Park, S.Y. Lee, H.J. Oh, M.A. Lim, W.T. Cho, E.K. Kim, J.H. Ju, Y.W. Park, S.H. Park, M.L. Cho, H.Y. Kim, TWEAK promotes the production of Interleukin-17 in rheumatoid arthritis, Cytokine 60(1) (2012) 143–9.

[59] T.S. Zheng, L.C. Burkly, No end in site: TWEAK/Fn14 activation and autoimmunity associated-end-organ pathologies, J Leukoc Biol 84(2) (2008) 338–47.

[60] L. Xia, H. Shen, W. Xiao, J. Lu, Increased serum TWEAK levels in Psoriatic arthritis: relationship with disease activity and matrix metalloproteinase-3 serum levels, Cytokine 53(3) (2011) 289–91.

[61] S. Orita, T. Koshi, T. Mitsuka, M. Miyagi, G. Inoue, G. Arai, T. Ishikawa, E. Hanaoka, K. Yamashita, M. Yamashita, Y. Eguchi, T. Toyone, K. Takahashi, S. Ohtori, Associations between proinflammatory cytokines in the synovial fluid and radiographic grading and pain-related scores in 47 consecutive patients with osteoarthritis of the knee, BMC musculoskeletal disorders 12 (2011) 144.

[62] D.H. Manicourt, P. Poilvache, A. Van Egeren, J.P. Devogelaer, M.E. Lenz, E.J. Thonar, Synovial fluid levels of tumor necrosis factor alpha and oncostatin M correlate with levels of markers of the degradation of crosslinked collagen and cartilage aggrecan in rheumatoid arthritis but not in osteoarthritis, Arthritis and rheumatism 43(2) (2000) 281–8.

[63] S. Rayego-Mateos, J.L. Morgado-Pascual, A.B. Sanz, A.M. Ramos, S. Eguchi, D. Batlle, J. Pato, G. Keri, J. Egido, A. Ortiz, M. Ruiz-Ortega, TWEAK transactivation of the epidermal growth factor receptor mediates renal inflammation, The Journal of pathology 231(4) (2013) 480–94.

[64] M. Skoumal, G. Kolarz, G. Haberhauer, W. Woloszczuk, G. Hawa, A. Klingler, Osteoprotegerin and the receptor activator of NF-kappa B ligand in the serum and synovial fluid. A comparison of patients with longstanding rheumatoid arthritis and osteoarthritis, Rheumatol Int 26(1) (2005) 63–9.

[65] L. Pulsatelli, P. Dolzani, T. Silvestri, P. Caraceni, A. Facchini, G. Ravaglia, C. Salvarani, R. Meliconi, E. Mariani, Soluble receptor activator of nuclear factor- kappaB Ligand (sRANKL)/osteoprotegerin balance in ageing and age-associated diseases, Biogerontology 5(2) (2004) 119–27.

[66] D.M. Findlay, G.J. Atkins, Osteoblast-chondrocyte interactions in osteoarthritis, Curr Osteoporos Rep 12(1) (2014) 127–34.

[67] D. Muratovic, D.M. Findlay, R.D. Quarrington, X. Cao, L.B. Solomon, G.J. Atkins, J.S. Kuliwaba, Elevated levels of active Transforming Growth Factor β1 in the subchondral bone relate spatially to cartilage loss and impaired bone quality in human knee osteoarthritis, Osteoarthritis Cartilage 30(6) (2022) 896–907.

[68] G.J. Atkins, P.S. Rowe, H.P. Lim, K.J. Welldon, R. Ormsby, A.R. Wijenayaka, L. Zelenchuk, A. Evdokiou, D.M. Findlay, Sclerostin is a locally acting regulator of late-osteoblast/preosteocyte differentiation and regulates mineralization through a MEPE-ASARM-dependent mechanism, J Bone Miner Res 26(7) (2011) 1425–36.

[69] A.R. Wijenayaka, M. Kogawa, H.P. Lim, L.F. Bonewald, D.M. Findlay, G.J. Atkins, Sclerostin stimulates osteocyte support of osteoclast activity by a RANKL-dependent pathway, PLoS One 6(10) (2011) e25900.

